# Identification of hsa-miR-106a-5p as an impact agent on promotion of multiple sclerosis using multi-step data analysis

**DOI:** 10.1101/2020.09.14.296483

**Authors:** Samira Rahimirad, Mohammad Navaderi, Shokoofeh Alaei, Mohammad Hossein Sanati

## Abstract

Multiple Sclerosis (MS) is a chronic, demyelinating disease in which the neuron myelin sheath is disrupted and leading to signal transductions disabilities. The evidence demonstrated that gene expression patterns and their related regulating factors are the most critical agents in Multiple Sclerosis demyelinating process. A miRNA is a small non-coding RNA which functions in post-transcriptional regulation of gene expression. Identification of specific miRNA dysregulation patterns in multiple sclerosis blood samples compared to healthy control can be used as a diagnostic and prognostic agent. Through the literature review and bioinformatics analysis, it was found that the hsa-miR-106a-5p can be considered as a significant MS pathogenic factor, which seems has an abnormal expression pattern in patients’ blood. Experimental validation using Real-Time PCR assay was carried to verifying the miR-106a-5p expression in Multiple Sclerosis and healthy control blood samples. The obtained results proved the miR-106a dysregulation in MS patients. The expression levels of miR-106a-5p were significantly down-regulated (Fold change=0.44) in patient blood samples compared to controls (*p*=0.059). Our study suggested that miR-106a-5p may have a biomarker potential to the diagnosis of MS patients based on its dysregulation patterns in Multiple Sclerosis blood.

## Introduction

Multiple sclerosis (MS) is a chronic, autoimmune, inflammatory disease of the central nervous system (CNS). Generally, MS pathogenicity involves the brain white matter and cause demyelination in the brain and spinal [1]. The myelin sheath insulates the nerves axons which cause fast transmission of the nerves signals through the neural cells. Accordingly, demyelination leading to discontinuous or retardation in CNS signal transmission [2]. Followed by demyelinating, axons may damage because of insulation loss and lack of nutrition feeding by oligodendrocytes. The inflammation affects oligodendrocytes activities through the MS progression [3].

About 2.5 million people are affected by MS around the world. The rate of MS is higher farther from the equator. MS is more common in women than men, the ratio may be as high as three to one [4]. There are four major types of MS including primary progressive multiple sclerosis (PPMS), relapsing-remitting multiple sclerosis (RRMS), secondary progressive multiple sclerosis (SPMS) and progressive relapsing multiple sclerosis (PRMS). Among 85 percent of patients, MS initiates with a relapsing-remitting. Approximately, 10-15 percent of patients suffer from PPMS which may gradually inter the PRMS phase [5]. MS symptoms include vision problems, muscle weakness and spasms, problems with balance and coordination, tingling, numbness, vertigo, dizziness and fatigue [6]. MS is a multi-factorial disorder caused by the interaction of genetic and environmental factors. Some genomic locus are known as MS susceptible factors like HLA-DR and HLA-DQ [7, 8]. The most effective environmental factors in pathogenicity of MS are the vitamin D deficiency, viral infections, geographic conditions and exposure to the sun [9].

MicroRNAs (miRNAs) are small non-coding RNAs (20 – 24 nucleotides) which regulate the mRNA stability and expression as the post-transcriptional modifications by targeting the 3’ untranslated region (UTR) of specific mRNA [10]. RNA polymerase II transcribes the miRNA genes, then the primary transcript related capping and polyadenylated processes are done and they fold as a stem-loop structure. The primary miRNA (Pri-miRNA) cleavages in the nucleus by the enzymatic activity of nuclear RNase III Drosha with the help of its essential cofactor DGCR8 [11]. After the cleavage step, the pre-miRNA is exported from the nucleus to the cytoplasm in association with Exportin-5 and Ran GTP. The pre-miRNA is cleaved by Dicer to a ~22-nt miRNA duplex. One of the two strands is assembled into the RNA-induced silencing complex (RISC) together with one of the Argonaute (Ago) proteins which facilitates the binding of mature miRNA to the 3’-untranslated region (UTR) of the target mRNA [12, 13]. MiRNA–mRNA complementarity causes a translational inhibition and/or degradation of the target mRNA [14]. MiRNAs play critical roles in a various biological process such as development, proliferation, differentiation, and inflammation. In the human immune system, miRNAs have an important role in modulating the immune response against the bacteria, virus, and other pathogens [15]. Also, miRNAs contribute to the biological activity of lymphocytes in the development and regulation of adaptive immunity [16]. Evidence has been shown that miRNA expression and function were associated with MS pathogenesis [17, 18]. As miRNAs are presence in biological fluids, such as blood, serum, plasma and cerebrospinal fluid (CSF), hence the circulating miRNAs are ideal prognostic factors for monitoring disease and response to treatment [19]. Recent studies have indicated the importance of the miR-106a in autoimmunity. Hsa-miR-106a (MI0000113) is a member of miR-106/363 cluster which comprises of the miR-106, miR-18b, miR-20b, miR-19b-2, miR-92a-2 and miR-363. These members are located in X chromosome [20].

In the present study, a multi-staged data mining followed by functional enrichment analysis was conducted and the expression levels of hsa-miR-106a-5p were examined by qRT-PCR. The main purpose of our study is to find potential miRNA and its targets involved in the pathogenesis of MS.

## Method

### Bioinformatics approach

#### Data collection

Literature mining along with Gene Expression Omnibus (GEO) datasets analysis was conducted to collect differential expressed (DE) miRNAs in MS blood samples. Our criteria were including the studies which (1) they analyzed the miRNA profile in blood samples of MS patients compared to control. (2) The patients were not taken any kinds of MS-related drugs. The animal model studies, cell line miRNAs profiling and studies of tissues were excluded. The datasets containing normalized data, GSE61741, GSE39644, GSE31568, GSE21079, and GSE17846 were analyzed to pick out the DE miRNAs using GEO2R tool (https://www.ncbi.nlm.nih.gov/geo/geo2r/) [21]. The GSE61741, GSE31568, and GSE17846 were based on Agilent <u>GPL9040</u> platform [febit Homo Sapiens miRBase 13.0] included 23, 23, 20 MS patients and 94, 70, 21 controls respectively. The GSE39644 was based on GLP15847 platform [NanoString nCounter Human miRNA assay] contains 8 blood samples of each patient and control, and the GSE21079 including 59 MS samples and 37 normal controls based on GPL8178 platform [Illumina Human v1 MicroRNA expression beadchip]. The overlapped DE miRNAs among these mentioned datasets were collected.

#### Functional Enrichment / miRNA-target interaction network Analysis

To found out the exact role of obtained miRNAs by data mining analysis, the functional enrichment survey was conducted using DIANA tools, miRPath v.3 [22]. The latest version of miRNA-target gene regulatory interaction networks (RegINs) file (miRTarBase release 6.1) were downloaded from CyTargetLinker and the networks were constructed and visualized using Cytoscape 3.4. [23].

#### RNA extraction and miRNA isolation procedure, Real-Time PCR

Using the Cochran formulate the sample size was calculated. Thirty-two Iranian MS patients from all clinical subtypes of MS who were diagnosed based on McDonald’s criteria [24,25] and 32 Iranian healthy individuals (control group) were enrolled for the study after signing an informed consent approved by the Ethics Committee. Sample collection was done under the supervision of the neurologist. Clinical characteristics of the individuals who are enrolled in this study shown in table 1. Total RNA was isolated from 500 μl whole blood samples using RNX-Plus kit (Cinna-gene, Iran) according to the manufacturer’s protocol. The isolated RNA quality and quantity was checked by NanoDrop instrument (Thermo Scientific™) and the absorbance at 260-280 nm was around 2 ng/ μl which is considered as a lack of DNA-protein contaminations or other inhibitors. Primers were designed based on miRNA sequences deposited in the miRBase database (version 20, June 2014) (http://www.mirbase.org/) with oligocalculator design program (http://biotools.nubic.northwestern.edu/OligoCalc.html) and primer blast software (https://www.ncbi.nlm.nih.gov/tools/primer-blast/). Using STAR-miR kit (NIGEB, Iran) the miRNA isolation and preparation of the reaction for Real-Time PCR was done according to the manufacturer’s protocol. In order to the miRNA purification, the poly-A polymerization was done based on kit protocol, the reaction contained 8μl on enzyme buffer, 1.6 μl dNTP mixture, 0.2 μl poly-A polymerase enzyme, 3 μl of extracted RNA in a 20 μl reaction volume. The reaction mixture was subjected to an initial incubation of 37°C for 60 min followed by 5 min at 85°C. The reverse transcription reactions were performed with the kit special adapter in 20-μL reactions following the manufacturer’s instructions.

**Table 1:** Clinical characteristics of the individuals enrolled in the study.

| MS<br>patient/Healthy | Male | Female | Age<br>Average | Treatment |
| --- | --- | --- | --- | --- |
| MS patients | 13 | 19 | 37.5 | 22 New case<br>10 non-treat |
| Healthy | 15 | 17 | 44.5 | - |

Real-time PCR was performed in triplicate with the Applied Biosystems StepOne™ (ABI) and each reaction contained 2.5 μl cDNA and 6.3 μl of SYBR^®^ Green (Ampliqon, Denmark) and 10 pmol of a mix of miRNA specific primers and housekeeping gene (5srRNA) primers with a total reaction volume of 12.5 μl. The reaction mixtures were subjected to initial denaturation of 95°C for 10 min followed by 38 cycles of 95°C for 30 sec, 59.4°C for 30 sec and 72°C for 30 sec. The melt curve analysis was performed by increasing the temperature from 65°C to 95°C with continuous 3% temperature increasing. The primers sequences used for the miRNA and the reference gene are given in Table 2. All quantitative PCR values were normalized to those of 5srRNA [26] as a housekeeping gene and the fold change of the target miRNA expression was calculated by the 2^−ΔΔCt^ formula [27]. All statistical analysis was done in SPSS version 24.

**Table 2:**
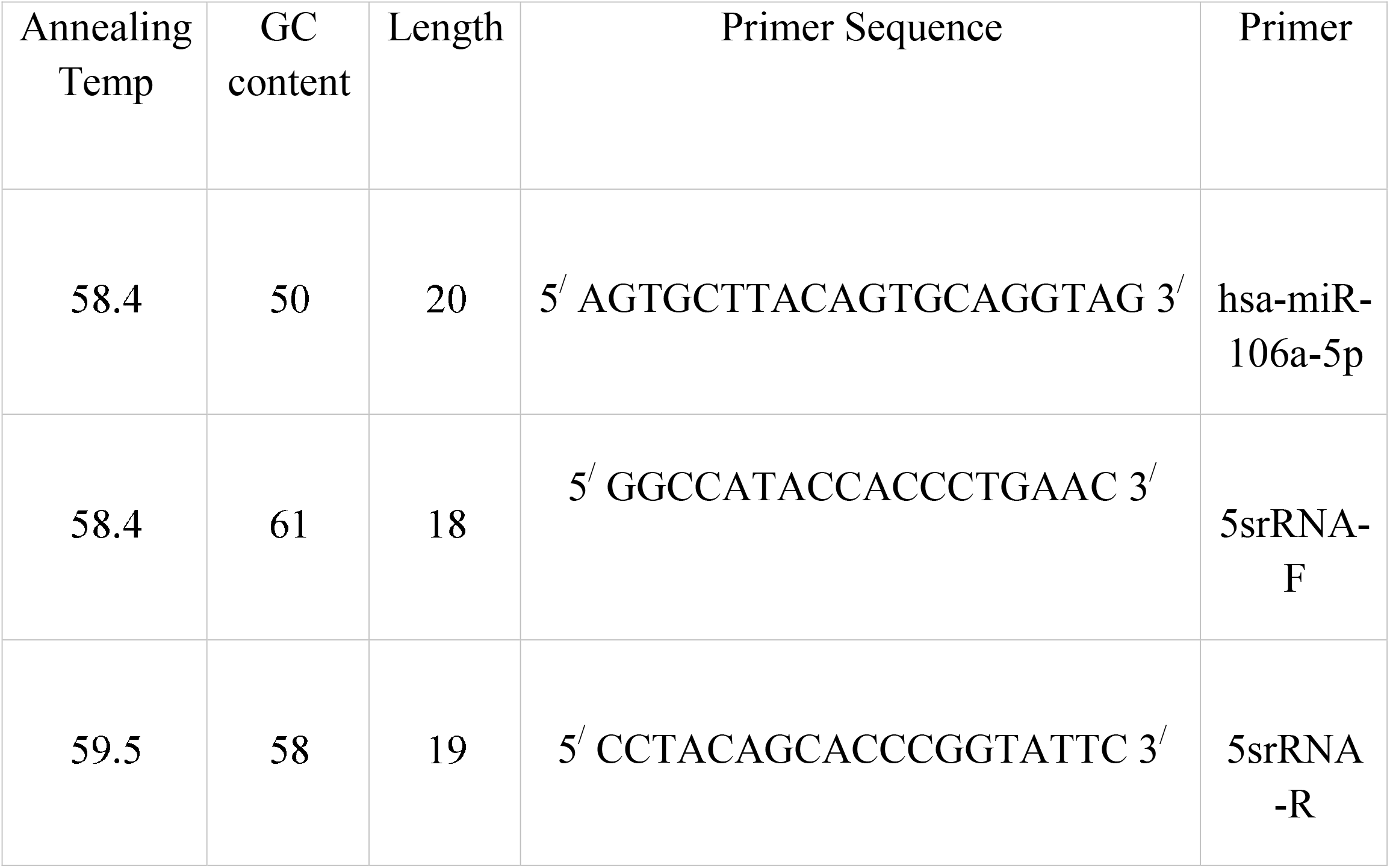
The primers sequence of hsa-miR-106a-5p and 5srRNA (housekeeping gene)

**Fig1.**
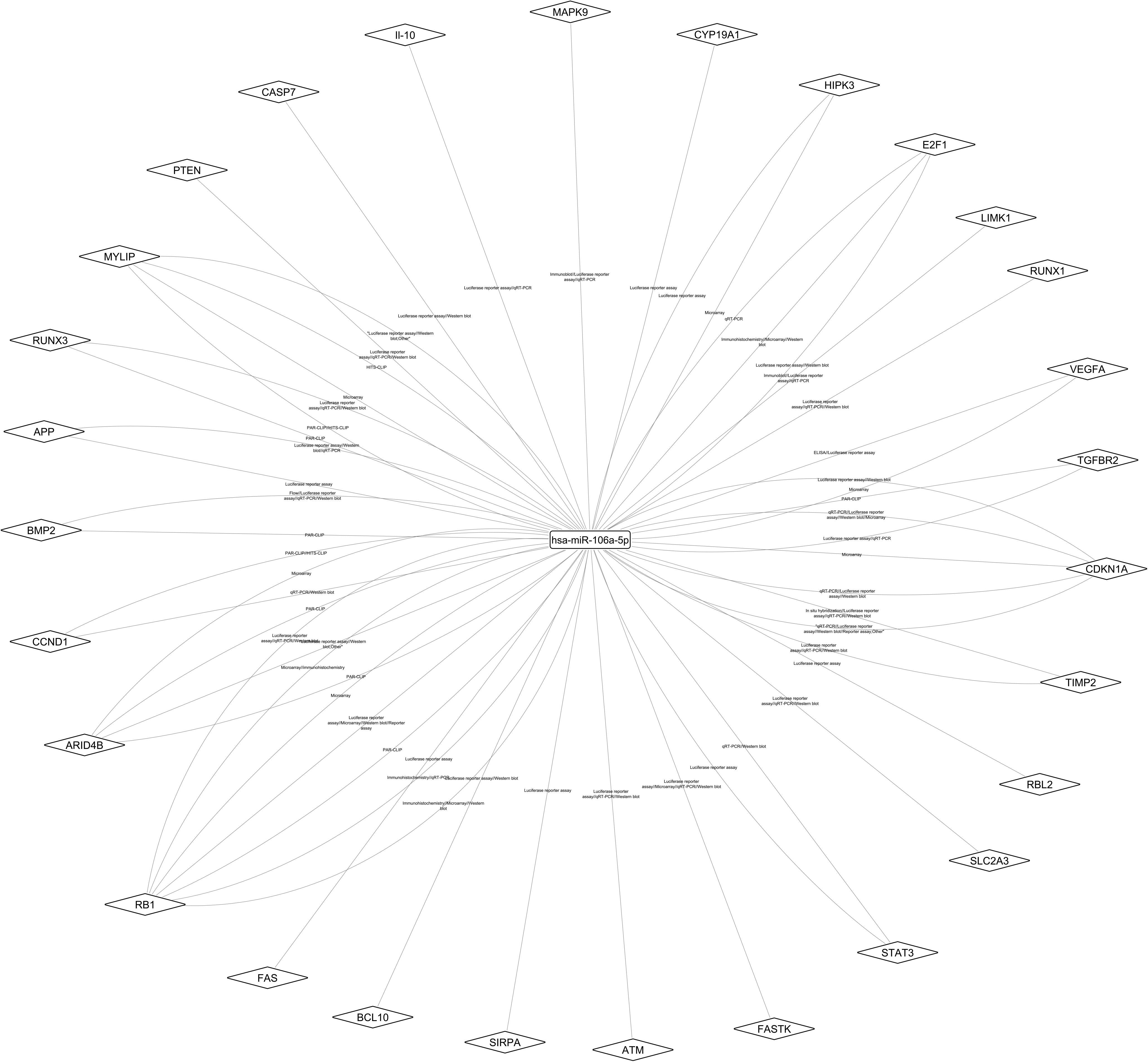
Schematic flow chart of the study.

## Result

### Selection of candidate miRNA in multiple Sclerosis pathogenesis using bioinformatics tools

Literature mining through MS-associated blood-derived microarray profiles revealed the dysregulated miRNAs. Based on our criteria, five studies including GSE61741, GSE39644, GSE31568, GSE21079, and GSE17846 were selected. From each dataset, the top 250 DE miRNAs were mined using GEO2R tool, then with the help of a Venn diagram, the overlapped miRNAs between datasets were collected. Figure 2 demonstrated that the 12 DE miRNAs comprise of hsa-miR-574-3p, hsa-miR-580, hsa-miR-30a, hsa-miR-146a, hsa-let-7g, hsa-let-7i, hsa-let-7b, hsa-miR-98, hsa-miR-106a, hsa-let-7c, hsa-miR-584, and hsa-miR-30e were similar among all five datasets. In table 3 the up/down regulation of each 12 miRNAs in datasets was documented.

**Fig2.**
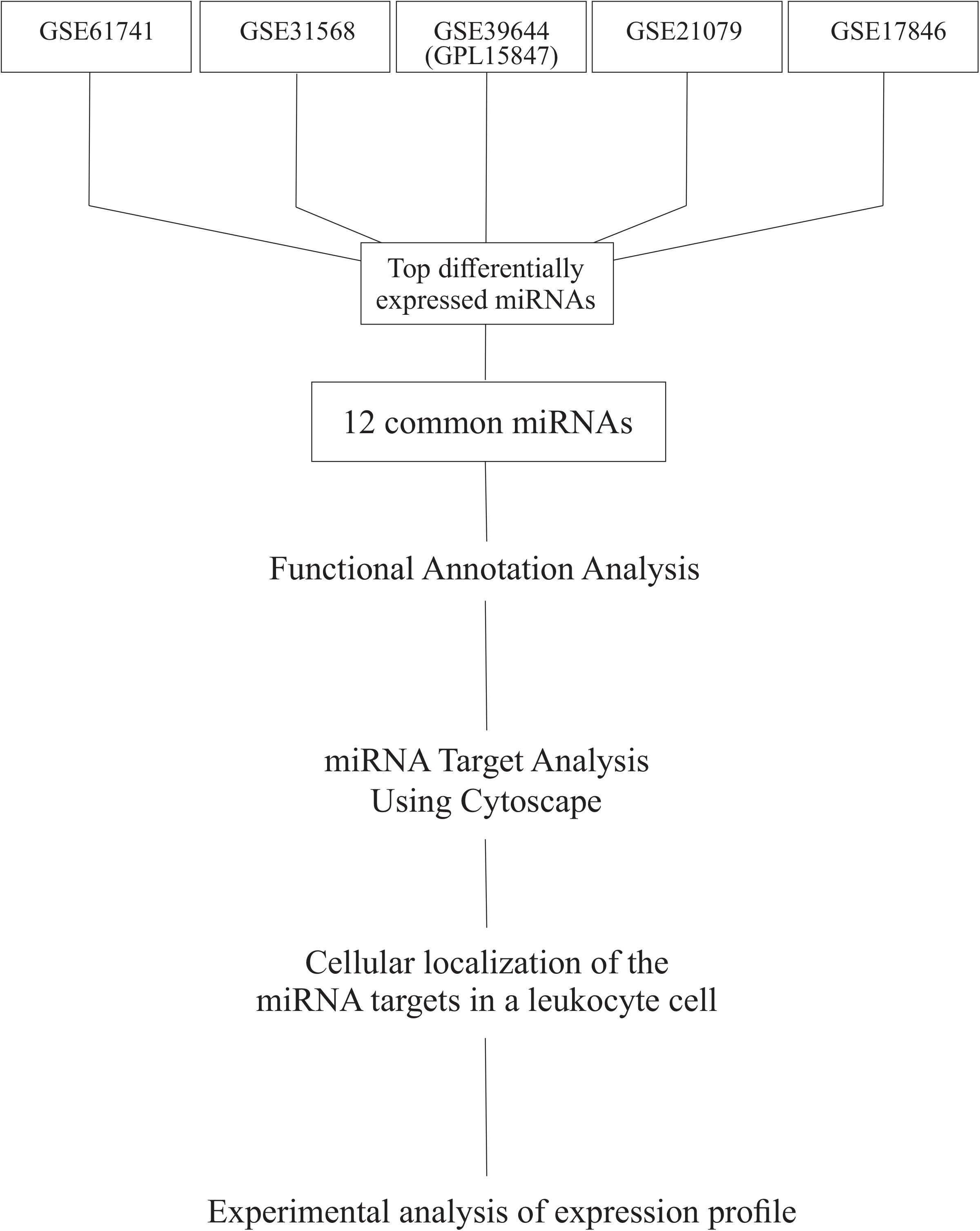
The Venn-diagram result of the overlapped miRNAs between datasets. In this analysis, we revealed that 12 miRNAs were showed differential expression between mentioned 5 MS blood-derived microarray datasets.

**Table 3:** The expression profile of obtained 12 miRNAs in each dataset. Using GEO2R tool the fold change (logFC) and p-value were calculated.

| MiRNA name | GSE code | P value | logFC |
| --- | --- | --- | --- |
| hsa-miR-574-3p | GSE61741 | 7.88E-11 | -1.498 |
|  | GSE39644<br>(GPL15847) | 1.56E-01 | 0.2826 |
|  | GSE31568 | 2.89E-09 | -1.368 |
|  | GSE21079 | 0.054242 | -0.0565 |
|  | GSE17846 | 1.37E-04 | -0.978 |
| hsa-miR-580 | GSE61741 | 2.39E-02 | -1.052 |
|  | GSE39644<br>(GPL15847) | 1.50E-02 | -0.5169 |
|  | GSE31568 | 5.06E-03 | -1.468 |
|  | GSE21079 | 0.203458 | -0.1205 |
|  | GSE17846 | 3.65E-04 | -0.741 |
| hsa-miR-30a | GSE61741 | 1.52E-04 | 0.929 |
|  | GSE39644<br>(GPL15847) | 3.20E-03 | 0.4208 |
|  | GSE31568 | 4.98E-03 | 0.829 |
|  | GSE21079 | 0.111407 | 0.2032 |
|  | GSE17846 | 2.39E-04 | 0.645 |
| hsa-miR-146a | GSE61741 | 1.50E-03 | 0.641 |
|  | GSE39644<br>(GPL15847) | 4.17E-05 | 1.6774 |
|  | GSE31568 | 1.36E-03 | 0.796 |
|  | GSE21079 | 0.081131 | 0.1198 |
|  | GSE17846 | 8.93E-05 | 0.643 |
| hsa-let-7g | GSE61741 | 8.83E-05 | 1.614 |
|  | GSE39644<br>(GPL15847) | 2.07E-02 | 0.4665 |
|  | GSE31568 | 4.30E-05 | 2.062 |
|  | GSE21079 | 0.005829 | 0.2556 |
|  | GSE17846 | 9.98E-04 | 1.151 |
| hsa-let-7i | GSE61741 | 3.75E-07 | 1.959 |
|  | GSE39644<br>(GPL15847) | 1.26E-01 | 0.3666 |
|  | GSE31568 | 9.33E-06 | 2.26 |
|  | GSE21079 | 0.003594 | -0.158 |
|  | GSE17846 | 1.01E-02 | 0.989 |
| hsa-let-7b | GSE61741 | 4.53E-08 | 2.018 |
|  | GSE39644<br>(GPL15847) | 8.46E-06 | 1.0974 |
|  | GSE31568 | 1.75E-06 | 2.193 |
|  | GSE21079 | 0.073397 | 0.0992 |
|  | GSE17846 | 2.28E-03 | 1.267 |
| hsa-miR-98 | GSE61741 | 2.49E-05 | 2.797 |
|  | GSE39644<br>(GPL15847) | 3.95E-01 | 0.1585 |
|  | GSE31568 | 9.18E-05 | 2.733 |
|  | GSE21079 | 3.08E-05 | -0.5383 |
|  | GSE17846 | 2.26E-02 | 0.898 |
| hsa-miR-106a | GSE61741 | 2.49E-02 | 0.39 |
|  | GSE39644<br>(GPL15847) | 5.58E-01 | 0.1133 |
|  | GSE31568 | 1.78E-02 | 0.987 |
|  | GSE21079 | 0.000119 | -0.3653 |
|  | GSE17846 | 1.48E-04 | -0.513 |
| hsa-let-7c | GSE61741 | 3.18E-08 | 2.159 |
|  | GSE39644<br>(GPL15847) | 2.25E-01 | -0.222 |
|  | GSE31568 | 1.31E-07 | 2.49 |
|  | GSE21079 | 0.139519 | -0.1008 |
|  | GSE17846 | 7.45E-05 | 1.488 |
| hsa-miR-584 | GSE61741 | 1.55E-02 | 1.21 |
|  | GSE39644<br>(GPL15847) | 1.50E-02 | -0.5299 |
|  | GSE31568 | 1.13E-03 | 1.856 |
|  | GSE21079 | 0.081188 | -0.1428 |
|  | GSE17846 | 4.25E-04 | 1.934 |
| hsa-miR-30e | GSE61741 | 4.86E-05 | 1.267 |
|  | GSE39644<br>(GPL15847) | 9.79E-02 | -0.3486 |
|  | GSE31568 | 1.88E-04 | 1.76 |
|  | GSE21079 | 0.176561 | -0.1158 |
|  | GSE17846 | 2.59E-05 | 0.886 |

In order to identify the candidate miRNAs which mentioned in all blood-derived microarray analyses, the functional annotations of these 12 miRNAs were checked through the collection of MS-related gene ontology (GO) terms including GO:0002376 (immune system process), GO:0002224 (toll-like receptor signaling pathway), GO:0045087 (innate immune response), GO:0032481 (positive regulation of type I interferon production), GO:0060337 (type I interferon signaling pathway), GO:0050900(leukocyte migration), GO:0019221 (cytokine-mediated signaling pathway), GO:0042590 (antigen processing and presentation of exogenous peptide antigen via MHC class I), GO:0002479 (antigen processing and presentation of exogenous peptide antigen via MHC class I, TAP-dependent), GO:0002474 (antigen processing and presentation of peptide antigen via MHC class I), GO:0032728 (positive regulation of interferon-beta production), and GO:0019886 (antigen processing and presentation of exogenous peptide antigen via MHC class II). The number of involvement of each miRNA in our intended GO biological process terms was calculated which shown in table 4. As a result, hsa-miR-30a, hsa-miR-146a, hsa-let-7g, hsa-let-7i, hsa-let-7b, hsa-miR-106a, and hsa-miR-30e involved in all 12 terms and entered in the next step of this study and others were excluded.

**Table 4:**
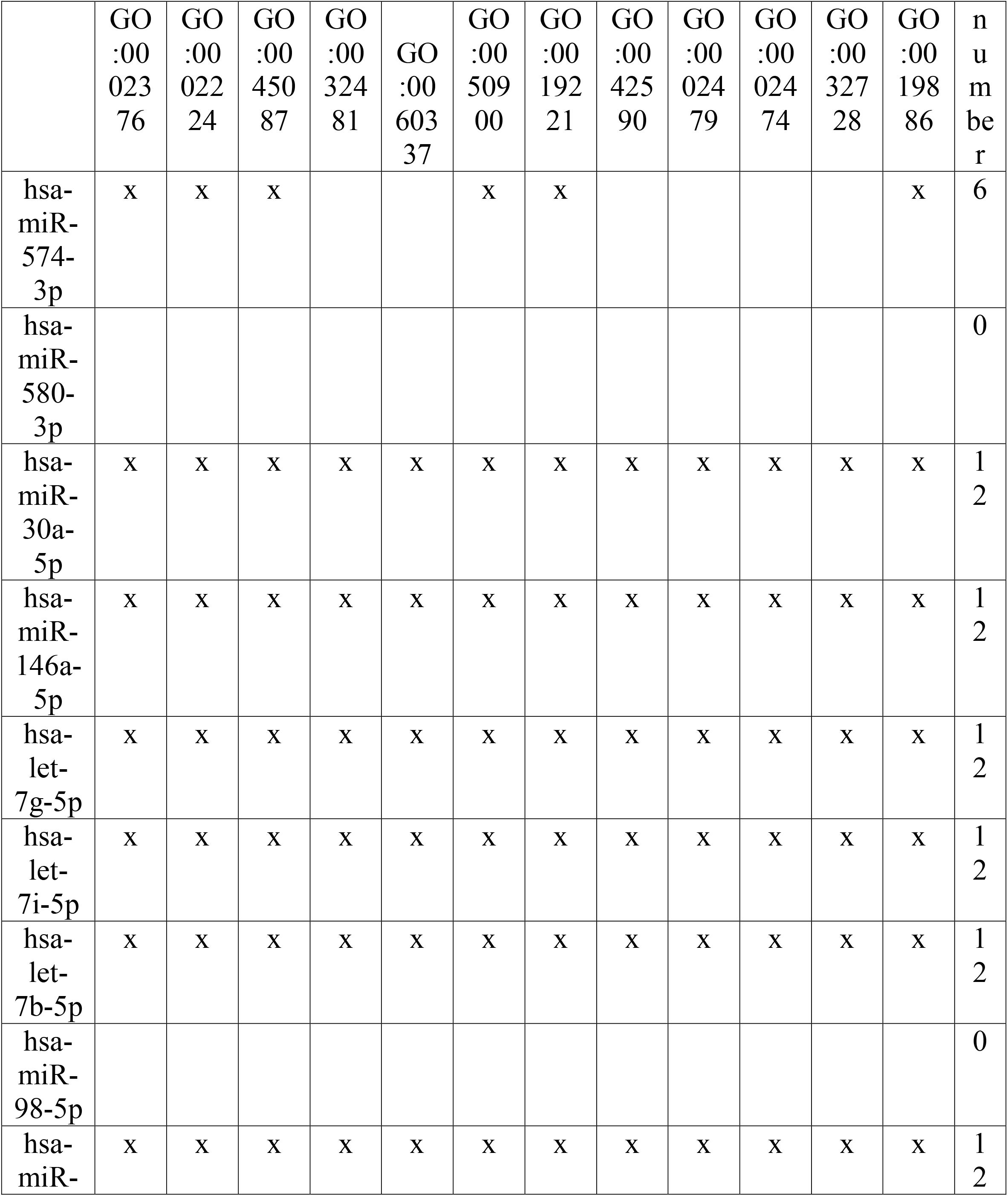
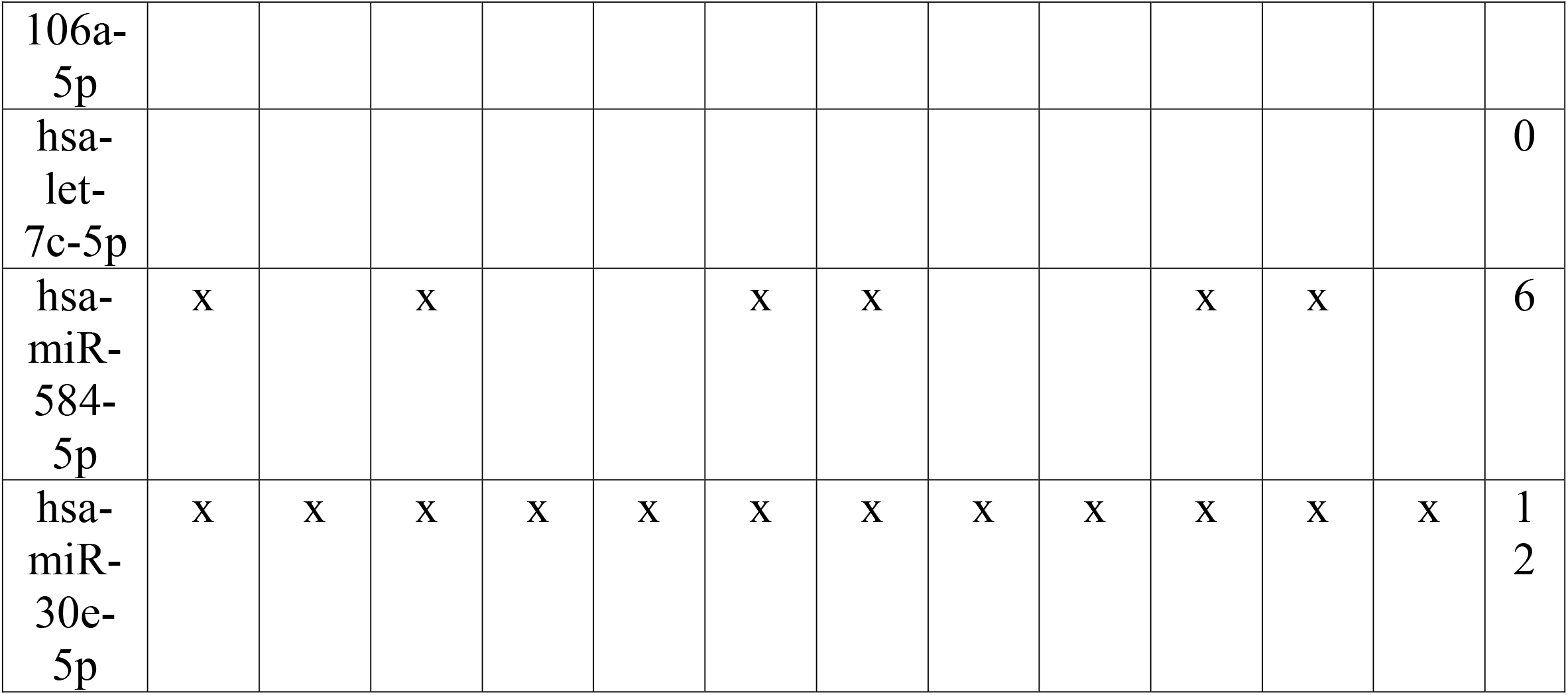
The involvement of collected miRNAs in MS-related GO terms. X indicates that the miRNA has a function in GO process.

Using the latest version of miRNA-target gene regulatory interaction networks (RegINs) file (miRTarBase release 6.1), the targets of 7 out of 12 miRNAs which impact on all GO process terms were obtained and visualized by constructing a network. This miRNA-target gene regulatory interaction network comprises of 2573 nodes which interacted by 18562 edges. Among these 7 miRNAs, hsa-let7-b-5p, hsa-miR-106a-5p, and hsa-miR-30a-5p have the most number of targets than the other ones. As this study aimed to identify the candidate miRNA which its dysregulation may play a significant role among the MS pathogenicity and there aren’t any studies that calculated the differential expression of the hsa-miR-106a-5p in blood samples of MS patients compared to controls using Real-Time PCR technique, hence mentioned miRNA was entered to detailed analysis. Among 668 targets of hsa-miR-106a-5p, only 28 genes have interacted with strong support as they are shown in figure 3. MS-related gene expression profile and functions of each 28 genes were retrieved through analysis of previous studies, then to visualized the impact of hsa-miR-106a-5p on its targets through the leukocyte cell different process, the pathway was constructed. For the first time in this pathway, we demonstrated MS-related processes in the leukocyte cell that involved genes are under expression regulation by hsa-miR-106a-5p. As shown in Figure 4, 28 strong target genes of hsa-miR-106a-5p are highlighted with white boxes which their functions such as transcription factor activity, Serine/Threonine kinase, Tumor inhibitor factor, Apoptosis inducer/inhibitor, Receptor, and ligand are distinct with different colors border. The inducing, inhibitory, phosphorylation, regulatory, and activator impacts of each gene on the other ones specified by color arrows which could show silencing activity of hsa-miR-106a-5p on one or more genes, may start the expression regulatory cascade.

**Fig3.**
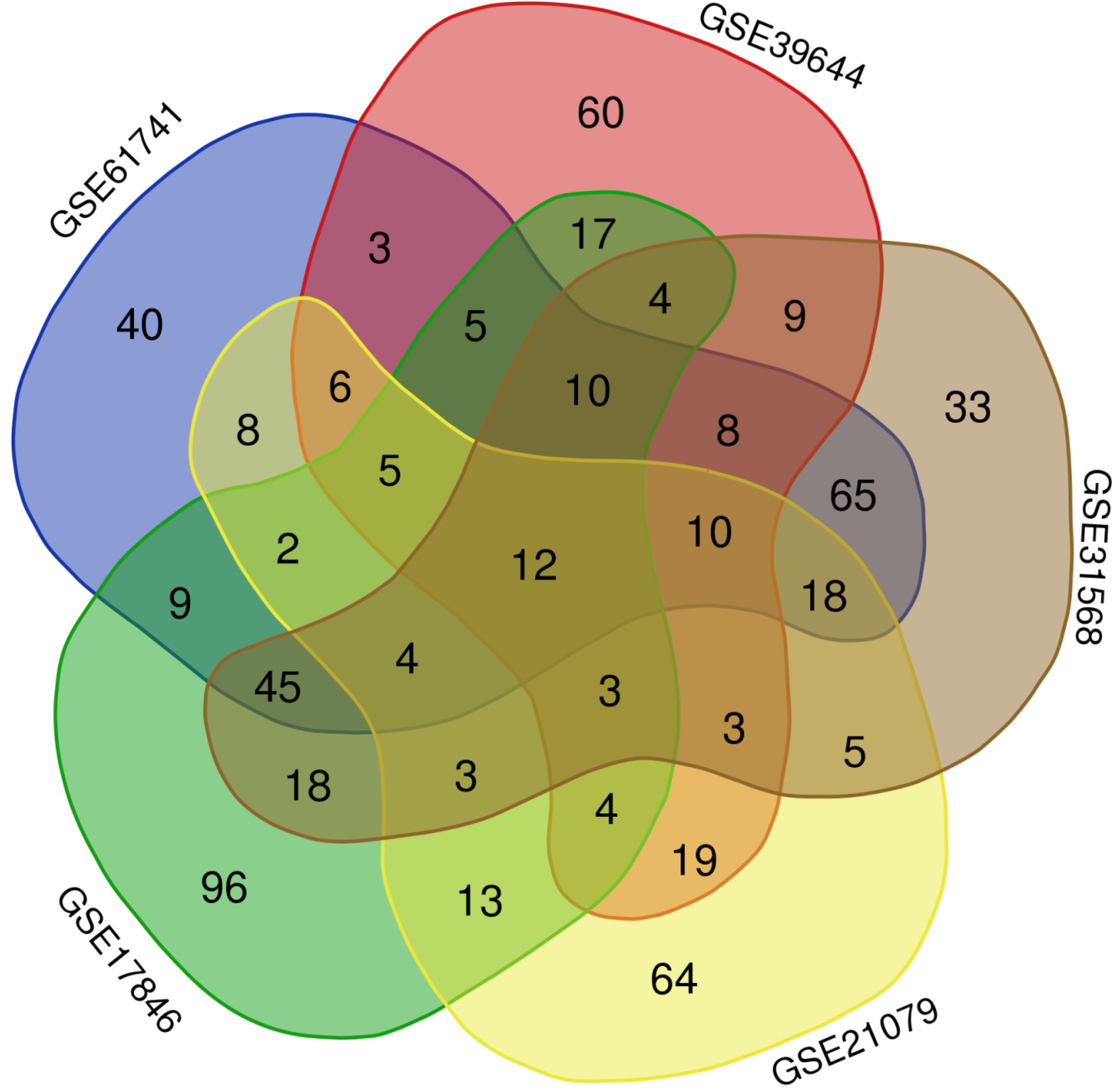
The miRNA-Target interaction network of hsa-miR-106a-5p. Using RegINs file, 668 gene targets were identified for hsa-miR-106a-5p that only 28 out of them interact with the strong support which in this figure are shown in diamond boxes. Edges are shown the experimental technique which validated the expression silencing role of hsa-miR-106a-5p on its target genes.

**Fig4.**
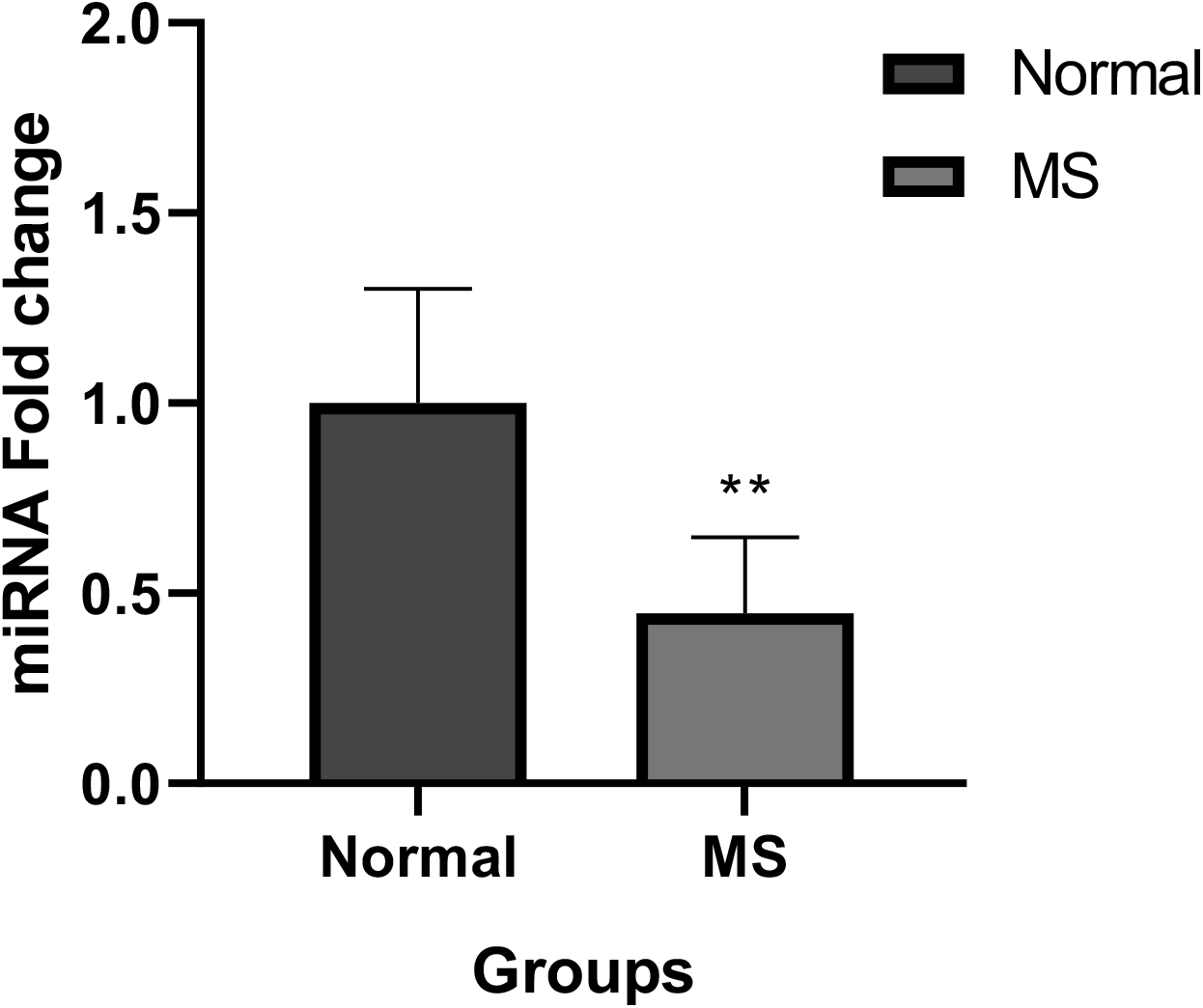
Cellular localization and functions of the targets of hsa-miR-106a-5p in a leukocyte cell. 28 target genes of hsa-miR-106a-5p are highlighted with white boxes and the other genes which are associated indirectly with these 28 genes are shown with grey boxes. The expression dysregulation of each gene is shown with an upper or down sign inside their boxes.

### Expression of hsa-miR-106a-5p in multiple sclerosis blood samples

The blood expression level of hsa-miR-106a-5p and 5srRNA as housekeeping gene was examined using qRT-PCR technique in 32 new case or non-treat patients and 32 healthy controls. The Ct values were collected by Applied Biosystems StepOne™ (ABI) software and the fold-change was calculated using the 2^−ΔΔCt^ formula. As a result of this expression examination, we demonstrated that the expression of hsa-miR-106a-5p was lower in blood samples of MS patients than in healthy controls (Fold-change= 0.447/ *P*<0/05) (Figure 5). analysis of 2^−ΔΔCt^ showed 0.44-fold decrease in levels of mentioned miRNA in the patient’s blood. ** represents p-value <0/05.

## Discussion

Multiple sclerosis is a chronic, inflammatory and neurodegenerative disease that leads to disabilities, vision problems and other neuromuscular impairments [1]. Today in general, the diagnostic method of this disease is examination of the demyelinated plaques presence in the central nervous system with the help of magnetic resonance imaging (MRI), which is not a completely specific diagnostic method [28]. So far, no method has been reported for early detection of this disease, but some have previously reported the presence of blood biomarkers in relation to the promotion of MS. Biomarkers are molecular markers that change in conditions of the disease in comparison with the natural conditions. MiRNAs are known as the most important biomarkers because the expression level of this group of small RNA molecules changes under different conditions and play a role in changing the profile of the expression of the genes in the body [29].

Therefore, finding a precise differential expression profile of a set of the most important pathogenic miRNAs in the blood of MS patients compared to healthy people may help to its early diagnosis, because any difference in blood levels of these factors may regulate the expression of various genes which could promote the various disorders such as MS. On the other hand, the diagnosis of a disease with these molecular factors can be more efficient and accurate than other methods [30]

In this study, bioinformatics analyses were conducted to find the most effective miRNAs in MS pathogenesis. For this purpose, miRNA expression studies in the blood of people with MS compared to healthy subjects which deposited into the Gene Expression Omnibus (GEO) database [31] were studied, including GSE61741, GSE39644, GSE31568, GSE21079, and GSE17846. The Venn Diagram was then mapped to find miRNAs that all the above studies confirmed their difference in expression. Eventually, 12 miRNAs were collected. Functional enrichment analysis along with identification of MS-related gene targets of candidate miRNAs leads to choose the hsa-miR-106a-5p as a potential factor in promoting the MS. The qRT-PCR technique was utilized to an examination of hsa-miR-106a-5p expression in blood samples of MS patients compared to control. Previously the decreased level of mentioned miRNA was measured by microarray [18] but there is no evidence which examines the blood level of hsa-miR-106a-5p in new case or non-treat patients. In this study, the blood level of hsa-miR-106a-5p was calculated as decreased 0.44-fold in patient samples than controls (*P*<0/05). Gene target interaction network of hsa-miR-106a-5p was constructed using miRTarBase file deposited in Cytoscape which resulted to obtain 28 targets with strong support interactions. The cellular localization and functions of each 28 target genes were demonstrated as a pathway (Figure 4) which shows how MS dysregulation of hsa-miR-106a-5p could impact on its targets expression profile and associated cell processes. Literature mining revealed that the level of *RBL2*, *APP*, *CYP19A1*, and *BMP2* increase during promotion of MS [32]. The study suggested that this increasing level may due to the MS-related downregulating of hsa-miR-106a-5p. In summary, the hsa-miR-106a-5p blood levels were decreased in all MS patients compared with the control group. This observation was remarkable and our results confirmed the previous reports.

## Conclusions

In all blood samples of MS compared with the control group, the hsa-miR-106a-5p levels were decreased. The decreased level of hsa-miR-106a-5p expression levels during MS promotion has been emphasized in numerous microarray studies. Using bioinformatics approaches we could identify the hsa-miR-106a-5p target genes which they function in different sites of a leukocyte cell and their dysregulated expression levels during the MS promotion may due to the impaired silencing activity of hsa-miR-106a-5p.

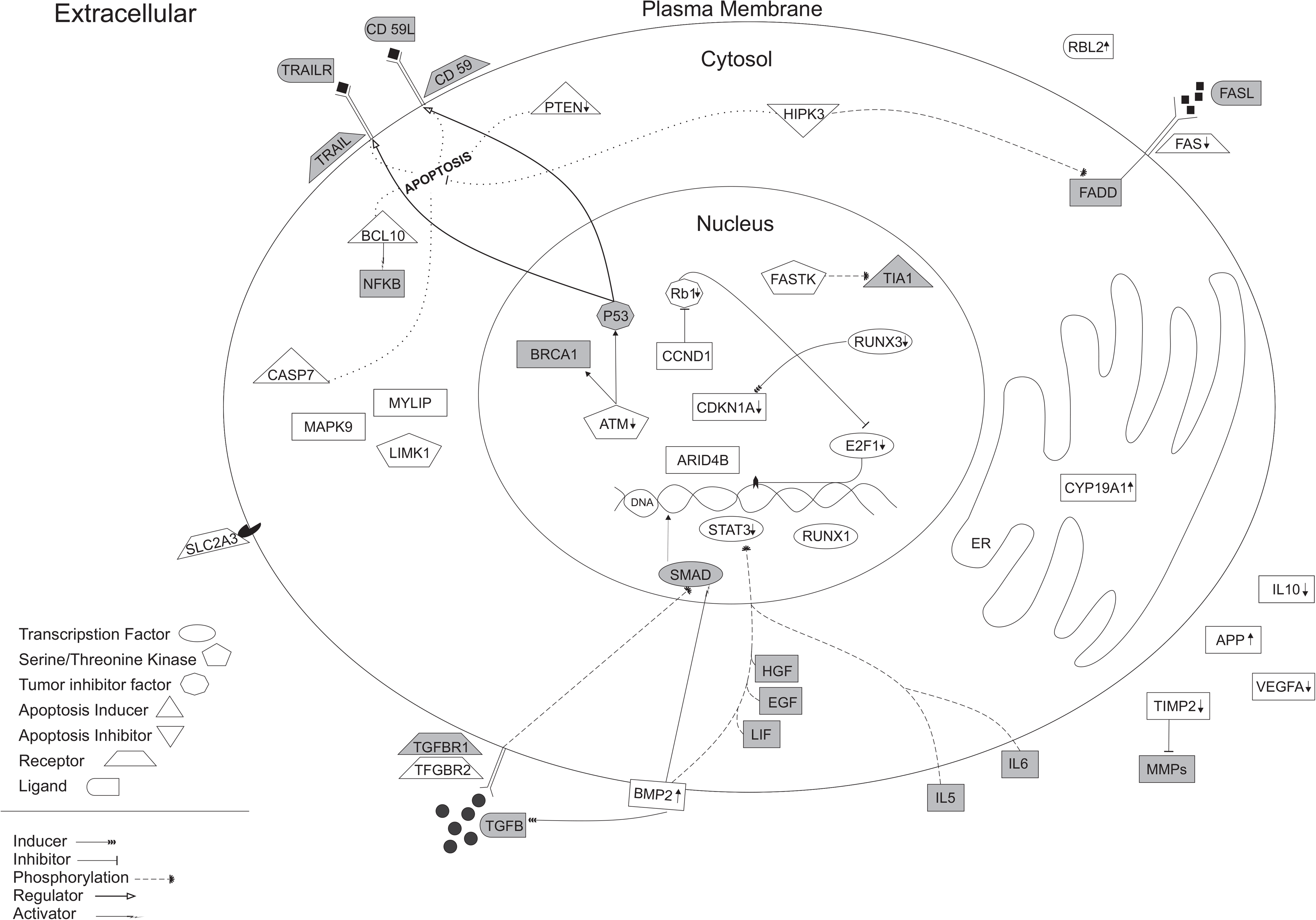

